# Inner-membrane transporter genotype shapes susceptibility to non-lytic antimicrobials and bacterial fitness in *Escherichia coli*

**DOI:** 10.64898/2026.09.04.749474

**Authors:** Elisabeth Dohmann Chang, Juliane Marie Jürgensen Engberg, Henrik Franzyk, Anders Løbner-Olesen, Jakob Frimodt-Møller

**Author notes:** Corresponding author: Jakob Frimodt-Møller.

## Abstract

Intracellularly acting antibacterials must cross the bacterial inner membrane to reach their targets. Using an isogenic *Escherichia coli* MG1655 panel carrying single and double deletions of *sbmA, ygdD* and *mdtM*, we assessed susceptibility to bleomycin, Oncocin, Api88 and Bac7(1–17), together with growth *in vitro* and competitive colonisation in the streptomycin-treated mouse intestine. SbmA was the dominant susceptibility determinant, whereas YgdD contributed most strongly to bleomycin and Api88 susceptibility. Deletion of *ygdD* or *sbmA* increased the bleomycin MIC four- and sixteenfold, respectively, while loss of both increased it 128-fold. Either single deletion also increased the Api88 MIC by more than 32-fold, whereas deletion of *mdtM* caused an eightfold increase. Single mutants grew similarly to wild type, whereas Δ*ygdD* Δ*sbmA* and Δ*ygdD* Δ*mdtM* had significantly longer doubling times; the largest effect was a 37% increase in Δ*ygdD* Δ*sbmA*. In mice, Δ*sbmA* and Δ*mdtM* showed reduced late competitive persistence. These findings support a compound-specific network of inner-membrane proteins whose fitness consequences depend on genetic background and environment.

## Introduction

The activity of an antibacterial compound with an intracellular target depends on both target engagement and intracellular exposure. In Gram-negative bacteria, compounds must traverse the outer membrane and the cytoplasmic membrane while avoiding degradation, sequestration and active efflux. For non-lytic antimicrobial peptides and bacteria-penetrating-peptide conjugates, transport across the cytoplasmic membrane can be the principal barrier to cytosolic delivery ^1^. Consequently, loss or alteration of an uptake pathway can reduce susceptibility without changing the intracellular target.

The compounds examined here comprise three linear proline-rich antimicrobial peptides (PrAMPs)— Oncocin, Api88 and Bac7(1–17)—that differ in primary sequence, length, net charge and terminal chemistry, together with bleomycin, a structurally distinct glycopeptide natural product with antibacterial activity and clinical use as an anticancer agent. Oncocin-derived peptides bind within the bacterial polypeptide-exit tunnel and inhibit protein synthesis ^2^, whereas Bac7 fragments interact with the ribosome and inhibit translation ^3^. Api88, an 18-residue analogue developed from apidaecin 1b, was initially reported to bind the chaperone DnaK ^4^; subsequent studies established the ribosome as a major target of apidaecin-derived PrAMPs, showing that Api137 traps release factors on terminating ribosomes ^5^, whereas C-terminally amidated Api88 occupies multiple sites within and near the polypeptide-exit tunnel and inhibits translation predominantly through a release-factor-independent, multimodal mechanism ^6^. Bleomycin differs fundamentally from these linear peptides: it contains a disaccharide-modified metal-binding domain connected through a methylvalerate–threonine linker to a bithiazole-containing DNA-binding tail, and its metal- and oxygen-dependent activation results in oxidative DNA strand cleavage ^7^. Thus, despite their distinct chemical architectures and intracellular mechanisms, the antibacterial activity of all four compounds requires access to targets within the bacterial cytoplasm, making efficient inner-membrane translocation critical.

SbmA is the best-characterised bacterial inner-membrane uptake system for PrAMPs. Genetic and uptake studies established that SbmA is required for efficient internalisation and full activity of several PrAMPs, including Bac7-derived peptides ^8,9^. Structural studies subsequently identified SbmA as a homodimeric, proton-coupled member of the SbmA-like peptide transporter family. An outward-open conformation was initially resolved ^10^, and more recent cryogenic electron microscopy structures captured occluded and inward-facing states consistent with proton-driven alternating access ^11^. These findings provide strong mechanistic support for direct SbmA-mediated transport into the cytosol.

Much less is known about YgdD, a poorly characterised inner-membrane protein. Loss of YgdD reduces susceptibility to arasin 1(1–25), but the available evidence does not distinguish direct peptide transport from an indirect contribution to uptake or intracellular activity ^12^. MdtM is a major facilitator superfamily transporter with established functions in multidrug and bile-salt efflux and monovalent-cation/proton antiport ^13–15^. Nevertheless, the *yjiL-mdtM* region has been implicated in the activity of selected PrAMP analogues, particularly when SbmA is absent ^16^. These observations suggest that susceptibility to intracellularly acting peptides may be determined by a network of inner-membrane proteins rather than by a single universal transporter.

Here, we examined all single and double combinations of *sbmA, ygdD* and *mdtM* deletions in *E. coli*. We asked whether the three proteins made compound-specific or overlapping contributions to susceptibility to bleomycin, Oncocin, Api88 and Bac7(1–17), and whether their loss affected growth in rich medium or the ability of the bacteria to colonise and persist competitively in the streptomycin-treated mouse intestine.

## Results

### Construction and verification of the transporter deletion panel

To determine the individual and combined contributions of SbmA, YgdD and MdtM to antimicrobial susceptibility and bacterial growth, an isogenic MG1655 deletion panel comprising the three single mutants and all three double mutants was assembled (Table 1). The complete panel was subsequently examined by MIC and growth analyses, while streptomycin-resistant derivatives of the three single mutants were used in competitive intestinal colonisation experiments.

**Table 1.** Susceptibility of the *E. coli* MG1655 transporter deletion panel to bleomycin and selected PrAMPs.

| Genotype | Bleomycin | Oncocin | Api88 | Bac7(1–17) |
| --- | --- | --- | --- | --- |
| Wild type | 0.5 | 15 | 1.9 | 32 |
| $\Delta sbmA$ | 8 | >60 | >60 | >64 |
| $\Delta ygdD$ | 2 | 30 | >60 | 64 |
| $\Delta mdtM$ | 0.5 | 15 | 15 | 32 |
| $\Delta sbmA \Delta mdtM$ | 8 | >60 | >60 | >64 |
| $\Delta ygdD \Delta sbmA$ | 64 | >60 | >60 | >64 |
| $\Delta ygdD \Delta mdtM$ | 8 | 60 | >60 | 32 |

MIC values are presented in $\mu\text{g/mL}$ . Values preceded by “>” indicate that growth inhibition was not achieved at the highest concentration tested.

### Transporter deletion profiles reveal compound-specific contributions of YgdD, SbmA and MdtM

The susceptibilities of the strain panel differed markedly among bleomycin, Oncocin, Api88 and Bac7(1–17) (Table 1). For bleomycin, the wild-type MIC was 0.5 µg/mL. Deletion of *ygdD* increased the MIC fourfold to 2 µg/mL, deletion of *sbmA* increased it sixteenfold to 8 µg/mL, and deletion of *mdtM* alone had no effect. The Δ*sbmA* Δ*mdtM* and Δ*ygdD* Δ*mdtM* mutants both had an MIC of 8 µg/mL, whereas the Δ*ygdD* Δ*sbmA* mutant had an MIC of 64 µg/mL. Thus, simultaneous loss of YgdD and SbmA was associated with a 128-fold MIC increase relative to wild type and a substantially greater resistance phenotype than loss of either protein alone. Api88 showed the strongest overall dependence on the tested proteins: the wild-type MIC was 1.9 µg/mL, deletion of *mdtM* increased it eightfold to 15 µg/mL, and the MIC exceeded 60 µg/mL in the Δ*ygdD* and Δ*sbmA* single mutants and in every double mutant containing either deletion. Oncocin and Bac7(1–17) showed a similar but less pronounced pattern dominated by SbmA. The Oncocin MIC increased from 15 µg/mL in wild type to 30 µg/mL in Δ*ygdD* and to >60 µg/mL in Δ*sbmA*; the Δ*ygdD* Δ*mdtM* strain had an MIC of 60 µg/mL, whereas all mutants containing Δ*sbmA* had MICs above 60 µg/mL. For Bac7(1–17), the MIC increased from 32 µg/mL in wild type to 64 µg/mL in Δ*ygdD* and to >64 µg/mL in Δ*sbmA*, whereas deletion of *mdtM*, either alone or together with *ygdD*, did not alter susceptibility. Overall, SbmA was the dominant susceptibility determinant, while the contributions of YgdD and MdtM varied with both the compound and the genetic background.

### Loss of YgdD together with SbmA or MdtM prolongs doubling time

None of the single-deletion mutants showed a detectable change in doubling time under the conditions tested. The Δ*sbmA*, Δ*mdtM* and Δ*ygdD* strains had mean doubling times of 24, 23 and 24 min, respectively, compared with 24 min for wild type. The Δ*sbmA* Δ*mdtM* mutant was also similar to wild type, with a mean doubling time of 24 min. In contrast, Δ*ygdD* Δ*sbmA* had a mean doubling time of 32 min, approximately 37% longer than wild type (P = 1.36 × 10^-6^). A more moderate but significant increase was observed for Δ*ygdD* Δ*mdtM*, which had a mean doubling time of 27 min (P = 0.02). The largest effect in Δ*ygdD* Δ*sbmA*, together with the absence of a detectable defect in either corresponding single mutant, is consistent with a genetic interaction and partially overlapping physiological functions. The smaller but significant effect in Δ*ygdD* Δ*mdtM* indicates that YgdD may also functionally interact with MdtM.

### Deletion of *sbmA* or *mdtM* reduces competitive persistence in the streptomycin-treated mouse intestine

To determine whether SbmA, YgdD or MdtM contributed to the ability of *E. coli* to colonise and persist competitively in the intestine, streptomycin-treated mice were inoculated with approximately equal numbers of wild type and the corresponding Δ*sbmA*, Δ*ygdD* or Δ*mdtM* mutant, and the two populations were monitored in faeces for 7 days (Fig. 1). On day 1, each mutant was recovered at a population size similar to that of its wild-type competitor, indicating that none of the deletions prevented initial establishment. In the Δ*sbmA* competition, the mutant and wild type remained at similar population sizes through day 3. Thereafter, the Δ*sbmA* population declined, while wild type remained near 6.5 log_10_ CFU/g faeces. On day 7, mean recovery was 4.48 log_10_ CFU/g for Δ*sbmA* and 6.57 log_10_ CFU/g for wild type, a difference of approximately 2.1 orders of magnitude; the original day-specific comparison yielded P = 0.03981. Variation among mice increased at the later time points, indicating that the extent of the decline differed among animals.

**Fig. 1.**
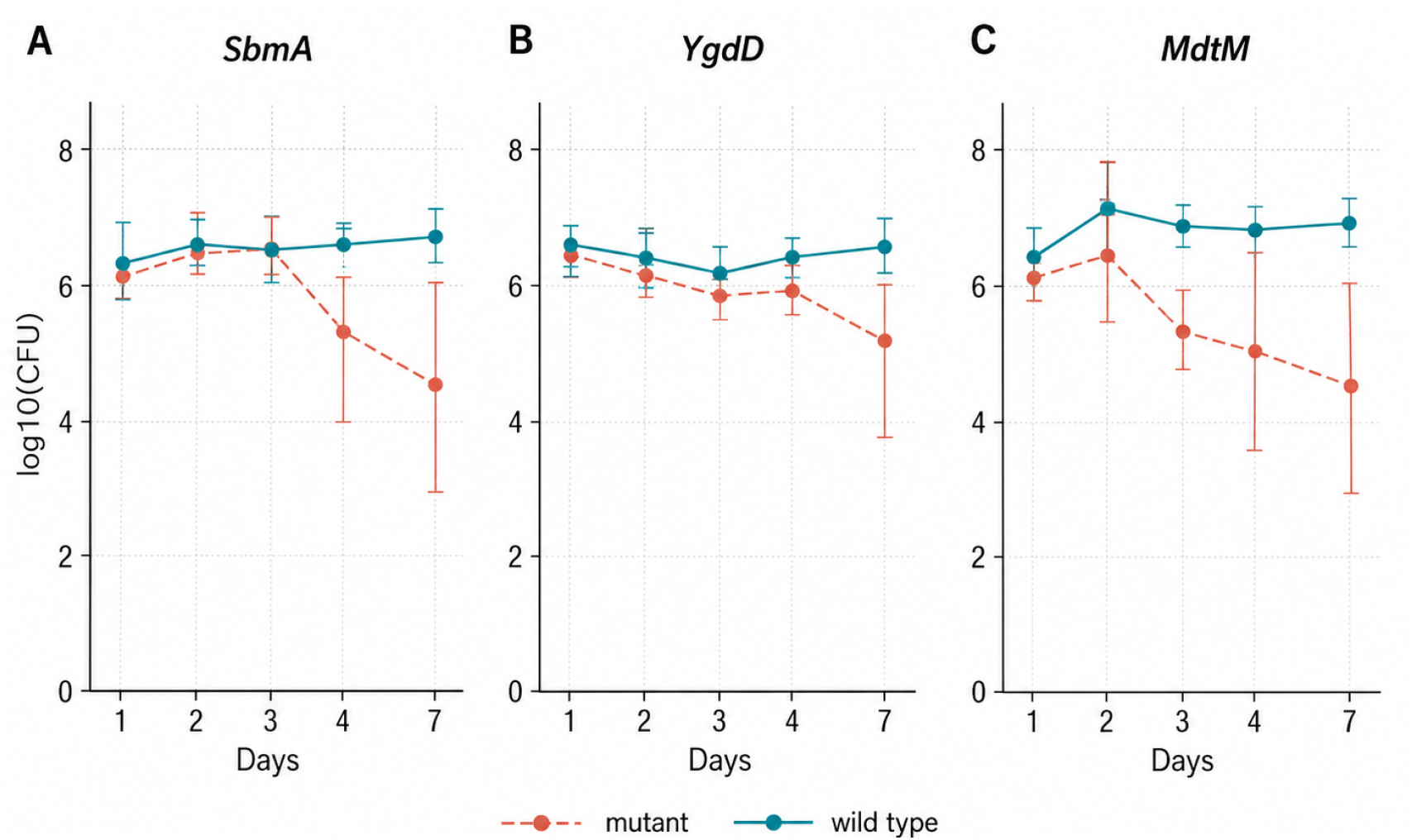
Competitive persistence of transporter deletion mutants in the streptomycin-treated mouse intestine. Streptomycin-treated mice were inoculated with approximately equal numbers of wild-type *E. coli* MG1655 and Δ*sbmA* (A), Δ*ygdD* (B) or Δ*mdtM* (C). Faecal samples were collected at the indicated times for 7 days, and the population sizes of wild type and mutant were estimated as described in Materials and methods. Points represent mean log_10_ CFU/g faeces, and error bars represent standard deviations. Five mice were assigned to each competition, although data were unavailable for some animals at individual time points. Where no mutant was identified among the 100 colonies examined, the mutant population was below the resolution of the colony-screening procedure and was not necessarily absent.

The Δ*ygdD* mutant showed a less pronounced phenotype. Wild type and Δ*ygdD* were recovered at comparable levels through day 4. By day 7, mean recovery of Δ*ygdD* had declined to 5.30 log_10_ CFU/g, compared with 6.54 log_10_ CFU/g for wild type. The original day-7 comparison was not significant (P = 0.1341), and the experiment therefore did not demonstrate a reproducible competitive disadvantage for Δ*ygdD* during the 7-day observation period.

The clearest reduction in competitive persistence was observed for Δ*mdtM*. On day 1, the mutant and wild type were recovered at 6.11 and 6.46 log_10_ CFU/g, respectively (P = 0.1599 in the original comparison). Wild type subsequently increased to approximately 7 log_10_ CFU/g and remained near that level, whereas Δ*mdtM* progressively declined after day 2. On day 3, mean recovery was 5.35 log_10_ CFU/g for Δ*mdtM* and 6.84 log_10_ CFU/g for wild type; the original comparison yielded P = 0.007181, although only three mice were represented at this time point. By day 7, mean recovery was 4.48 log_10_ CFU/g for the mutant and 6.95 log_10_ CFU/g for wild type (P = 0.02299). No Δ*mdtM* colonies were identified among the 100 colonies examined in three of the five day-7 samples, indicating that the mutant proportion was below the resolution of the colony-screening procedure rather than proving complete elimination.

## Discussion

This study shows that susceptibility to non-lytic, intracellularly acting compounds is shaped by a network of inner-membrane proteins rather than by SbmA alone, and that the contribution of individual proteins varies among compounds. YgdD contributed strongly to bleomycin and Api88 susceptibility and more modestly to Oncocin and Bac7(1– 17). For bleomycin, simultaneous loss of YgdD and SbmA increased the MIC substantially beyond either single deletion, while the same double mutant also showed the largest growth defect. Deletion of *sbmA* or *mdtM* was furthermore associated with reduced late competitive persistence in the streptomycin-treated mouse intestine. These findings extend the transporter-dependent intracellular-exposure framework described in our mini-review ^1^ by linking compound susceptibility to the physiological and environment-dependent consequences of losing the same inner-membrane proteins.

SbmA-dependent internalisation has been demonstrated genetically and directly for several PrAMPs, including Bac7-derived peptides ^8,9^, and structural studies support a proton-coupled alternating-access mechanism ^10,11^. The large SbmA-associated MIC shifts observed for all four compounds are therefore consistent with its established role in uptake. The present data extend this framework by showing that the contribution of the other inner-membrane proteins to cytosolic uptake is compound- and genotype-wdependent (Fig. 2).

**Fig. 2.**
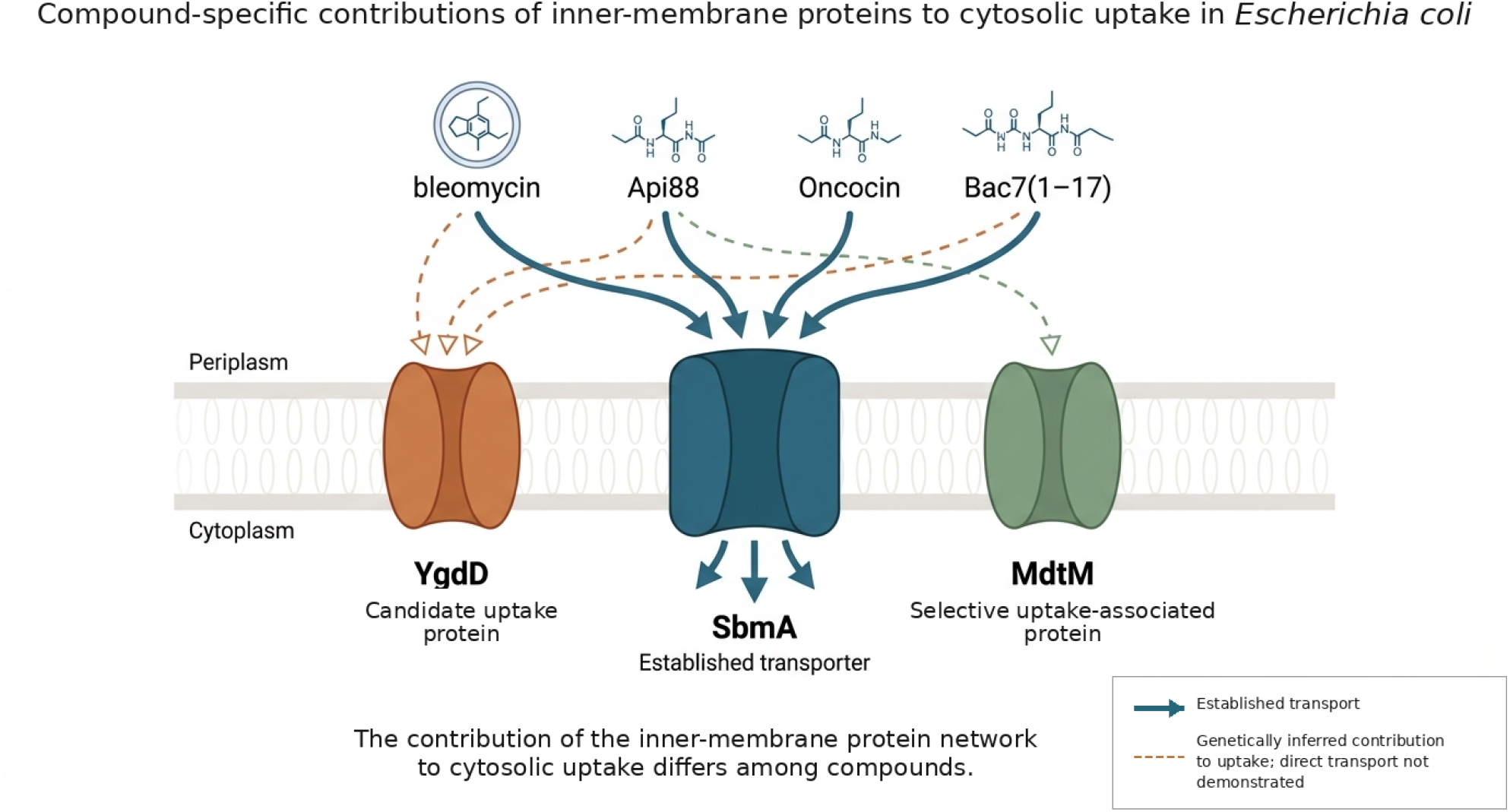
Conceptual model of compound-specific contributions of inner-membrane proteins to antimicrobial uptake in *Escherichia coli*. SbmA is shown as an established inner-membrane transporter contributing to the uptake of bleomycin, Api88, Oncocin and Bac7(1–17). The present genetic data implicate YgdD as an additional candidate uptake protein for all four compounds, with the strongest effects observed for bleomycin and Api88, whereas MdtM was selectively associated with Api88 uptake in the single-mutant background. Solid arrows indicate transport supported by published functional evidence. Dashed connectors indicate genetically inferred contributions to uptake for which direct transport has not been demonstrated. The inner-membrane protein network is shared, but the contribution of individual proteins to cytosolic uptake differs among compounds.

The clearest evidence for a previously unrecognised contribution of YgdD to antimicrobial susceptibility came from bleomycin. Bleomycin differs structurally and mechanistically from the PrAMPs examined here and ultimately damages DNA rather than inhibiting protein synthesis. Loss of SbmA has previously been associated with reduced bleomycin susceptibility in *Salmonella enterica* ^17^; the present work extends this observation to *E. coli* and identifies YgdD as an additional contributor. The highest level of bleomycin resistance was observed when both SbmA and YgdD were absent, which is consistent with separate or only partially overlapping contributions and argues against a model in which the YgdD phenotype is simply secondary to loss of SbmA activity. A similar, although less pronounced, pattern was observed for Oncocin and Bac7(1–17): the MICs increased primarily following loss of SbmA, as previously reported ^8,9^, while deletion of *ygdD* produced a twofold increase for each peptide. Because MIC measurements do not directly assess intracellular accumulation, these results identify YgdD as a susceptibility determinant and candidate uptake protein rather than demonstrating direct transport. Testing a broader panel of structurally diverse compounds could help define which molecular features determine YgdD dependence.

Although the Δ*sbmA* and Δ*ygdD* mutants showed high-level Api88 resistance, neither single mutant had a detectable growth defect *in vitro*. This resembles the absence of an obvious laboratory fitness cost in *S. enterica sbmA* mutants selected for PR-39 resistance ^17^ and illustrates how transporter-loss resistance may arise without an immediate penalty detectable by routine growth assays. Significant growth defects emerged, however, when *ygdD* was deleted together with *sbmA* or *mdtM*. The largest effect occurred in Δ*ygdD* Δ*sbmA*, suggesting that YgdD and SbmA make partially overlapping contributions to normal cellular physiology, while the smaller Δ*ygdD* Δ*mdtM* phenotype indicates an additional functional interaction involving MdtM. Because no antimicrobial compound was present during the growth experiments, these phenotypes reflect endogenous physiological functions rather than impaired uptake of the compounds tested. These genetic interactions should be investigated using independently reconstructed and complemented strains.

Deletion of *mdtM* alone produced a comparatively narrow susceptibility phenotype, increasing the MIC only for Api88. However, the Δ*ygdD* Δ*mdtM* mutant had higher bleomycin and Oncocin MICs than either corresponding single mutant, indicating that an MdtM-associated contribution can become apparent in the absence of YgdD. The *yjiL-mdtM* region has previously been implicated in the activity of selected apidaecin and oncocin analogues, particularly in an *sbmA*-deficient background ^16^. Consistent with a relationship between transporter genotype and intra- cellular exposure, label-free uptake measurements showed that transporter-deficient *E. coli* accumulated lower intracellular amounts of PrAMPs than wild type, with reduced peptide accumulation generally corresponding to increased MICs ^18^. Differences between the present and earlier findings may reflect peptide sequence, strain background, medium or assay conditions. Together, these observations reinforce that the contribution of individual uptake proteins should be defined for each compound and genetic background rather than assigned uniformly to an entire PrAMP family.

The intestinal competition experiments further suggest that the consequences of transporter loss depend on the environment. The Δ*sbmA* and Δ*mdtM* mutants established initially at levels comparable to wild type but declined during the later stages of the experiment, despite neither single mutant showing a growth defect in rich medium. The Δ*mdtM* phenotype is biologically plausible given the established roles of MdtM in alkaline pH homeostasis through monovalent-cation/proton antiport and in protection from bile-salt stress ^14,15^. The intestinal environment exposes bacteria to changing pH, cation concentrations and bile salts, providing a possible explanation for normal growth of Δ*mdtM* in MHBI but reduced competitive persistence in mice. The Δ*ygdD* mouse phenotype remained inconclusive. Thus, the combination of high-level Api88 resistance, no detectable single-mutant growth defect and no reproducible 7-day intestinal defect identifies loss of YgdD as a potentially low-cost resistance route under the conditions tested, although larger and appropriately powered competition experiments are needed.

In conclusion, the results support a shift from a predominantly SbmA-centred view of inner-membrane uptake to a model in which a shared set of inner-membrane proteins contributes differently to the cytosolic entry of individual compounds. This has practical implications for the development of non-lytic antimicrobials. Testing candidates only against wild type measures the combined effects of target inhibition and cellular entry, while testing only a Δ*sbmA* mutant may miss secondary uptake proteins and genetic interactions among pathways. Profiling candidate compounds against a small panel of uptake, envelope and membrane-energetics mutants would help identify agents that depend excessively on a single non-essential entry route. Compounds that retain activity when one route is absent, either through autonomous uptake or through multiple non-redundant pathways, may be less vulnerable to transporter-loss resistance.

## Materials and methods

### Bacterial strains

The *E. coli* strains used in this study are listed in Table 3, and the oligonucleotides used for strain construction are listed in Table 4. The previously described streptomycin-resistant MG1655 strain ^19^ served as the parental strain. The *sbmA* and *ygdD* deletion/replacement alleles were obtained from the Keio collection ^20^ and introduced into MG1655 by P1 transduction ^21^ to generate Δ*sbmA* and Δ*ygdD*, respectively. The chromosomal region spanning *yjiL-mdtM* was deleted by λ Red recombineering ^22^. A chloramphenicol-resistance cassette was amplified from pKD3 using primers yjiL_pKD3_FW and mdtM_pKD3_RV and introduced into wild-type cells carrying pKD46. Chloramphenicol-resistant transformants were selected, and the deletion was verified by PCR to generate Δ*yjiL-mdtM* (hereafter Δ*mdtM*).

**Table 2.**
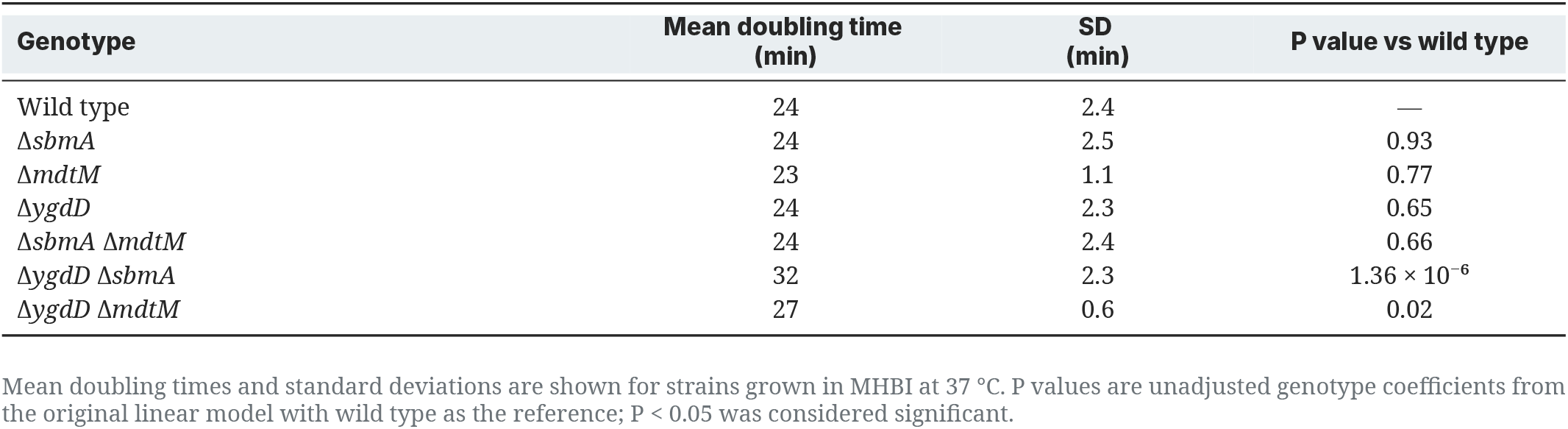
Doubling times for wild type and the single- and double-deletion mutants.

**Table 3.** List of strains.

| Relevant genotype | Abbreviated form | Source/reference |
| --- | --- | --- |
| MG1655; F- λ-rph-1 streptomycin resistant | Wild type | 19 |
| $\Delta sbmA^a$ $Kan^R$ | $\Delta sbmA$ | This study |
| $\Delta ygdD^a$ $Kan^R$ | $\Delta ygdD$ | This study |
| $\Delta yjiL\text{-}mdtM^a$ $CM^R$ | $\Delta mdtM$ | This study |
| $\Delta ygdD \Delta sbmA^a$ | $\Delta ygdD \Delta sbmA$ | This study |
| $\Delta mdtM \Delta sbmA^a$ | $\Delta mdtM \Delta sbmA$ | This study |
| $\Delta ygdD \Delta mdtM^a$ | $\Delta ygdD \Delta mdtM$ | This study |

<sup>a</sup> Otherwise as wild type.

**Table 4.**
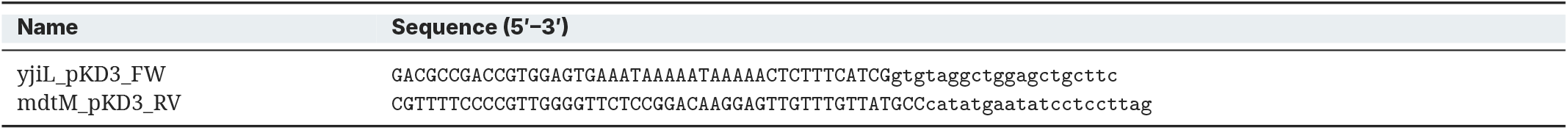
Oligonucleotides and genome-editing reagents.

For construction of multiple-deletion strains, resistance cassettes were removed from the relevant recipient backgrounds using pCP20 as described previously ^22^. The *sbmA* deletion/replacement allele was introduced by P1 transduction into Δ*ygdD* and Δ*mdtM* to generate Δ*ygdD* Δ*sbmA* and Δ*mdtM* Δ*sbmA*, respectively. Similarly, the *ygdD* deletion/replacement allele was introduced into Δ*mdtM* to generate Δ*ygdD* Δ*mdtM*. All strains were verified by PCR.

### Minimum inhibitory concentration determination

Minimum inhibitory concentrations (MICs) were determined by broth microdilution using a modification of standard procedures ^23–25^. Briefly, an overnight bacterial culture was diluted to approximately 5 × 10^5^ CFU/mL in MHBI. Bacterial suspension (100 µL) was dispensed into a low-binding 96- well microtitre plate (Thermo Scientific, catalogue no. 260895) together with 11 µL of test compound prepared as a twofold serial dilution. Vehicle-control wells contained 100 µL bacterial suspension and 11 µL of the vehicle used to prepare the corresponding test compound. Plates were incubated without shaking for 18–24 h at 37 °C unless otherwise stated. The MIC was defined as the lowest concentration that prevented visible growth, corresponding to OD_595_ < 0.1. Inoculum density was verified by viable counting in representative experiments. MICs for all test compounds were determined in three biological replicates.

### Growth measurements and doubling-time calculation

Doubling times were determined from three independent cultures grown in MHB II at 37 °C with aeration. Overnight culture (10 µL) was inoculated into 50 mL medium in a 250-mL flask. OD_595_ was measured during exponential growth, between approximately OD 0.05 and 0.5, at 5–10-min intervals. The specific growth rate (µ) was obtained from the slope of ln(OD) versus time over the exponential phase, and doubling time was calculated as ln(2)/µ.

### Streptomycin-treated mouse competition experiments

The streptomycin-treated mouse model used to compare the large-intestinal colonising abilities of *E. coli* strains has been described previously ^26,27^. Briefly, six- to eight-week-old outbred female NMRI (Naval Medical Research Institute) mice were given drinking water containing streptomycin sulfate (5 g/L) for 24 h to reduce the resident facultative anaerobic microbiota ^28^. Mice were orally inoculated with 100 µL of 20% (wt/vol) sucrose containing 10^6^ CFU of LB-grown *E. coli*. Faecal population sizes were used to estimate colonisation of the large intestine ^26^. Faeces were collected 24 h after inoculation and at the subsequent indicated time points for up to 7 days. Mice were housed in groups of five and individually marked to allow collection of faecal pellets from each animal. Fresh drinking water containing streptomycin sulfate (5 g/L) was provided daily. Each faecal sample was homogenised in 1% Bacto tryptone (Difco Laboratories, NJ, USA), diluted in the same medium and plated on MacConkey agar containing the appropriate antibiotics. When required, 1 mL of faecal homogenate was centrifuged at 12,000 × g, resuspended in 100 µL of 1% Bacto tryptone and plated, increasing the detection sensitivity from 10^2^ to 10 CFU/g faeces. Strains were distinguished by plating on lactose MacConkey agar containing streptomycin alone or in combination with kanamycin or chloramphenicol. Plates were incubated for 18–24 h at 37 °C. When necessary, 100 colonies from streptomycin-containing plates were tooth-picked onto plates containing the appropriate additional antibiotic. All animal experiments were approved by the Animal Experiments Inspectorate, Danish Ministry of Justice (permit no. 2007/561-1430), and were performed by trained personnel.

### Statistical analysis

Doubling-time data were analysed using a linear model with genotype as a categorical predictor and wild type as the reference. Mouse population sizes were log_10_-transformed and compared between wild type and mutant at selected time points using two-tailed Welch’s t-tests. Because wild type and mutant were recovered from the same animals and the mice were sampled repeatedly, the day-specific mouse comparisons are considered exploratory. P < 0.05 was considered significant; the reported P values were not adjusted for multiple comparisons.

